# Repurposing antihypertensive drugs for the prevention of Alzheimer’s disease: a Mendelian Randomization study

**DOI:** 10.1101/486878

**Authors:** Venexia M Walker, Patrick G Kehoe, Richard M Martin, Neil M Davies

**Affiliations:** MRC University of Bristol Integrative Epidemiology Unit, Bristol, UK; Bristol Medical School: Population Health Sciences, University of Bristol, Bristol, UK; Dementia Research Group, Bristol Medical School, University of Bristol, Bristol, UK; Bristol Medical School: Translational Health Sciences, University of Bristol, Bristol, UK

## Abstract

**Background:** Evidence concerning the potential repurposing of antihypertensives for Alzheimer’s disease prevention is inconclusive. We used Mendelian randomization, which can be more robust to confounding by indication and patient characteristics, to investigate the effects of lowering systolic blood pressure (SBP), via different antihypertensive drug classes, on Alzheimer’s disease.

**Methods:** We used summary statistics from genome wide association studies of SBP (from UK Biobank) and Alzheimer’s disease (from the International Genomics of Alzheimer’s Project) in a two-sample Mendelian randomization analysis. We identified single nucleotide polymorphisms (SNPs) that mimic the action of antihypertensive targets and estimated the effect of lowering SBP, via antihypertensive drug classes, on Alzheimer’s disease. We also report the effect of lowering SBP on Alzheimer’s disease by combining all drug targets and without consideration of the associated drugs.

**Results:** There was limited evidence that lowering SBP, via antihypertensive drug classes, affected Alzheimer’s disease risk. For example, *calcium channel blockers* had an odds ratio (OR) per 10mmHg lower SBP of 1.53 (95% confidence interval (CI): 0.94 to 2.49; p=0.09; SNPs=17). We also found limited evidence for an effect of lowering SBP on Alzheimer’s disease when combining all drug targets (OR per 10mmHg lower SBP: 1.14; 95%CI: 0.83 to 1.56; p=0.41; SNPs=59) and without consideration of the associated drug targets (OR per 10mmHg lower SBP: 1.04; 95%CI: 0.95 to 1.13; p=0.45; SNPs=153).

**Conclusions:** Lowering SBP itself is unlikely to affect risk of developing Alzheimer’s disease. Consequently, if specific antihypertensive drug classes do affect risk of Alzheimer’s disease, they are unlikely to do so via SBP.

**KEY MESSAGES:**

- This is the first study to use Mendelian randomization to estimate the effects of the twelve most common antihypertensive drug classes on Alzheimer’s disease.
- Lowering systolic blood pressure itself is unlikely to affect risk of developing Alzheimer’s disease.
- If specific antihypertensive drug classes do affect Alzheimer’s disease risk, they are unlikely to do so via systolic blood pressure.

## INTRODUCTION

Drug repurposing applies existing drugs to novel indications to identify potential treatments in a more rapid and cost-effective manner than traditional drug development. This approach is of interest for Alzheimer’s disease as there are currently no preventative or disease-modifying therapies, despite investment in 1120 unique drug targets between 1995 and 2014. (1–3) Antihypertensive drugs have previously been highlighted as priority repurposing candidates for Alzheimer’s disease prevention and several observational studies and a handful of trials have investigated this hypothesis. (2,4) However, the evidence to date is inconclusive.

Mendelian randomization has been proposed to predict drug repurposing opportunities and overcome some of the issues associated with conventional observational studies. (5) Mendelian randomization is a form of instrumental variable analysis that uses germline genetic variation, assigned randomly at conception and akin to randomization in a randomized controlled trial, as an instrument for potentially modifiable exposures of interest. (6–8) Without individual level data, two-sample Mendelian randomization can be implemented using summary data on single nucleotide polymorphisms (SNPs) from separate genome wide association studies (GWAS) for the instrument-exposure (sample one) and instrument-outcome (sample two) associations. (9) This approach has been used before to study the relationship between blood pressure and Alzheimer’s disease but it has not been used to estimate the effects of the twelve most common antihypertensive drug classes on Alzheimer’s disease. (10–12)

In this study, we use SNPs as instruments, selected to mimic the action of the protein targets of antihypertensive drug classes, in a two-sample Mendelian randomization analysis of systolic blood pressure on Alzheimer’s disease. Our rationale is to understand if there are differences between specific antihypertensive drug classes on Alzheimer’s disease risk, which could inform the prioritization of repurposing candidates, and provide evidence at the drug class level that could be triangulated with that from other sources. (13) Greater understanding of antihypertensives and their effect on Alzheimer’s disease may also highlight potentially relevant biological mechanisms for this disease. Some of these drugs, such as those acting through angiotensin receptor and calcium channel blocking mechanisms, have been suggested to have protective effects on Alzheimer’s disease that are independent of blood pressure lowering. (14–16) As we used instruments that proxy the protein targets, our estimates include all downstream effects of altering these targets, regardless of whether they are a direct result of lowering systolic blood pressure. (5)

## METHODS

### Study design

We conducted a two-sample Mendelian randomization analysis using summary data on SNPs from GWAS. We identified SNPs to proxy exposure to an antihypertensive drug on the basis that they mimicked the action of that drug on their molecular targets. For example, angiotensin-converting enzyme inhibitors work by inhibiting the enzyme angiotensin-converting enzyme. We therefore selected SNPs in the angiotensin-converting enzyme gene to use as a genetic proxy for this drug class. Effect sizes for these SNPs were then extracted from a GWAS of systolic blood pressure to estimate the instrument-exposure association. (17) The instrument-outcome association was estimated using the effect sizes for these same SNPs from a GWAS of Alzheimer’s disease. (18) All data used were publicly available and mostly obtained from European ancestry populations.

### Systolic blood pressure phenotype

The systolic blood pressure phenotype was defined using a GWAS of the UK Biobank cohort. (17) UK Biobank consists of 503,317 Caucasian people from the UK, aged between 38 years and 73 years. (19,20) The GWAS was based on 317,754 of the participants.

### Alzheimer’s disease phenotype

The Alzheimer’s disease phenotype was defined using the International Genomics of Alzheimer’s Project (IGAP) GWAS Stage 1 results. (47). These data were from a meta-analysis of 17,008 Alzheimer’s disease cases and 37,154 controls of European ancestry. (48).

### Instrument selection

We identified twelve antihypertensive drug classes in the British National Formulary. (21) They were: *adrenergic neurone blocking drugs; alpha-adrenoceptor blockers; angiotensin-converting enzyme inhibitors; angiotensin-II receptor blockers; beta-adrenoceptor blockers; calcium channel blockers; centrally acting antihypertensive drugs; loop diuretics; potassium-sparing diuretics and aldosterone antagonists; renin inhibitors; thiazides and related diuretics;* and *vasodilator antihypertensives*. Using the drug substance information, we were able to identify pharmacologically active protein targets and the corresponding genes in the DrugBank database (https://www.drugbank.ca/; version 5.1.1). (22) We then identified SNPs to instrument each target using the Genotype-Tissue Expression (GTEx) project data (Release V7; dbGaP Accession phs000424.v7.p2), which contains expression quantitative trait loci analysis of 48 tissues in 620 donors. (23) The full GTEx dataset, which consists of 714 donors, is 65.8% male and 85.2% white. SNPs marked as the ‘best SNP’ for the gene (defined by GTEx as the variant with the smallest nominal p-value for a variant-gene pair) in any tissue were selected for analysis.

To validate the SNPs as instruments for antihypertensive drug targets, we estimated their effect on systolic blood pressure using two-sample Mendelian randomization. The SNP-expression association, extracted from GTEx as described above, was on the scale of a standard deviation change in RNA expression levels for each additional effect allele. The SNP-systolic blood pressure association was extracted from the systolic blood pressure GWAS in UK Biobank and represented the standard deviation change in systolic blood pressure for each additional effect allele. These associations were then used to estimate the effect of the protein target on systolic blood pressure (i.e. the standard deviation change in systolic blood pressure per standard deviation change in RNA expression levels). SNPs with evidence of an effect on systolic blood pressure were retained for the main analysis. This instrument selection process is presented in Supplementary Figure 1.

### Statistical methods

We used two-sample Mendelian randomization to estimate the effect of lowering systolic blood pressure on Alzheimer’s disease in three ways. First, we estimated the effect of specific drug classes by combining the effects of any of the drug targets associated with a given drug class. This used the instruments defined in the previous section. Second, we estimated the effect of antihypertensive drugs as a whole on Alzheimer’s disease by combining all drug targets. Again, this used the instruments defined in the previous section. Finally, we estimated the overall effect of systolic blood pressure on Alzheimer’s disease by combining the effects of any genome-wide significant SNPs for systolic blood pressure.

When multiple SNPs were being used as an instrument, ‘clumping’ was performed to identify independent SNPs using the linkage disequilibrium between them. SNPs absent in the outcome data were replaced by proxy SNPs in high linkage disequilibrium from the 1000 Genomes Project European data where possible. (24,25) Proxies were required to have a minimum R-squared value of 0.8 and palindromic SNPs were permitted if their minor allele frequency was less than 0.3.

Prior to the analysis, data were harmonised to represent an increase in systolic blood pressure. Mendelian randomization was then performed using the inverse variance weighted method or, for single-SNP instruments, the Wald ratio. (26–28) Once complete, the Mendelian randomization results were transformed to be the odds ratio (OR) for Alzheimer’s disease per 10mmHg lower systolic blood pressure to make the effect comparable to taking an antihypertensive, which on average reduces systolic blood pressure by 9mmHg. (29) All analyses used genome reference consortium human build 37 (GRCh37), assembly Hg19, and were performed in R using the package ‘TwoSampleMR’. (24)

### Sensitivity analyses

Mendelian randomization estimates may be subject to horizontal pleiotropy, whereby the SNP(s) chosen to proxy the exposure affect the outcome by a different mechanism to that intended. (30) To estimate the extent of horizontal pleiotropy, we applied MR-Egger regression to all estimates based on ten or more SNPs. The regression intercept for these analyses “can be interpreted as an estimate of the average pleiotropic effect across the genetic variants”. (31) This can detect directional pleiotropy, which occurs when the biasing effects are not balanced around the null.

To examine heterogeneity within the drug classes, we also considered the effects of individual drug targets on Alzheimer’s disease. This analysis allowed us to ascertain whether certain targets were driving the drug class effects we observed. Drug classes with very heterogeneous target results can be considered to have less reliable estimates than those where targets were more homogeneous.

### Code availability

The analysis used R version 3.4.4. (32) All coding files are available from GitHub (https://github.com/venexia/MR-antihypertensives-AD).

## RESULTS

### Instrument selection

We identified a total of 73 unique protein targets of antihypertensive drugs (Supplementary Table 1). Among these targets, 68 had an effect in one or more GTEx tissues and 58 of those 68 provided evidence that the target affected systolic blood pressure (Supplementary Table 2). Supplementary Figure 2 summarizes the results of the Mendelian randomization analysis of expression on systolic blood pressure. A further six targets were excluded prior to the main analysis because neither the genetic instrument, nor a suitable proxy, were available in the outcome GWAS. Consequently, 52 unique protein targets were ultimately analysed (Supplementary Table 3).

**Figure 1:**
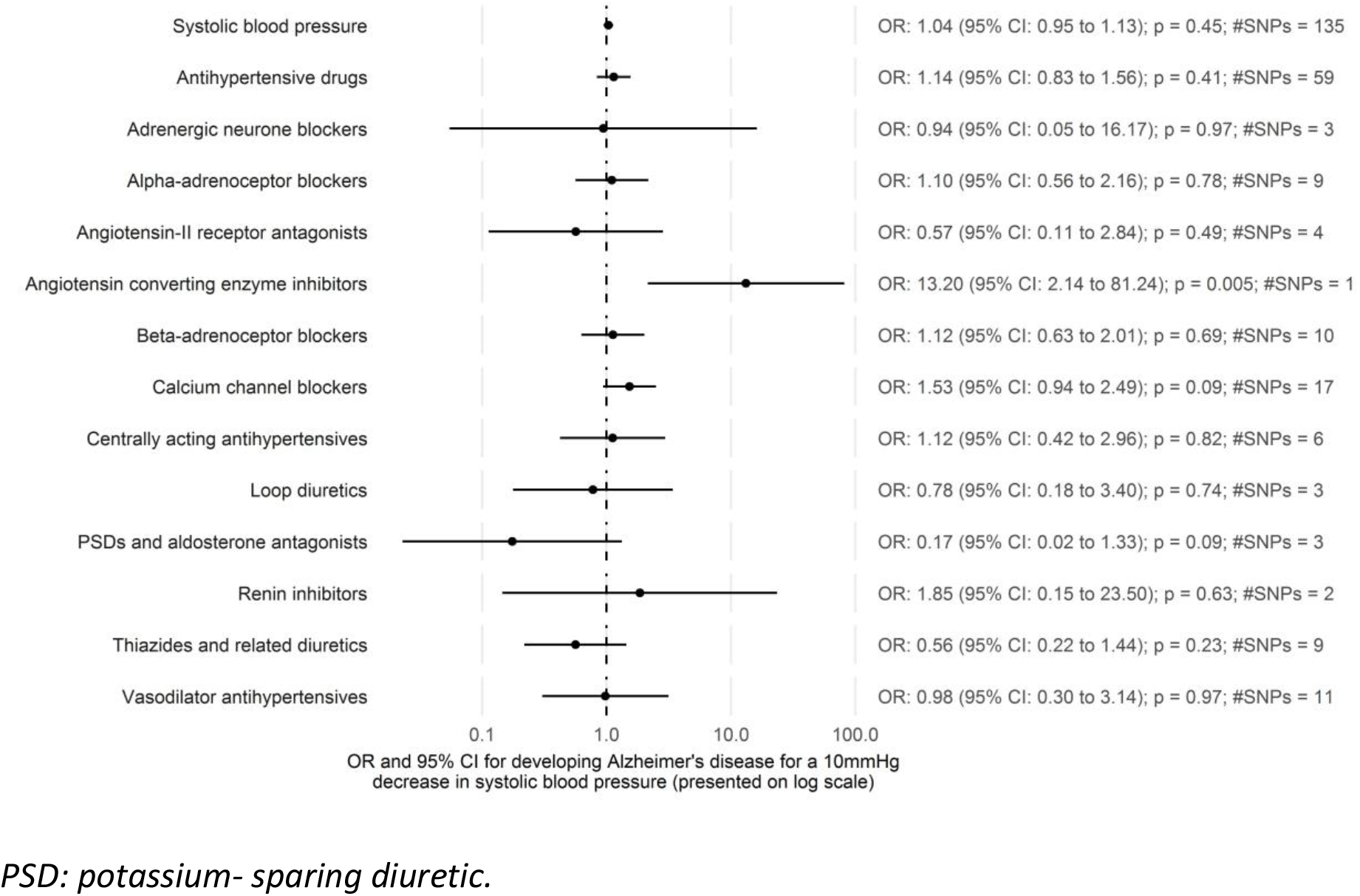
Estimates for the effect of systolic blood pressure on Alzheimer’s disease from two-sample Mendelian randomization

### Drug class effects

There was limited evidence that reducing systolic blood pressure affected risk of Alzheimer’s disease at the drug class level with most estimates failing to exclude the null (Supplementary Table 4). For example, *calcium channel blockers* had an OR of 1.53 (95% CI: 0.94 to 2.49; p=0.09; SNPs=17) and *loop diuretics* an OR of 0.78 (95% CI: 0.18 to 3.40; p=0.74; SNPs=3) per 10mmHg lower systolic blood pressure. The exceptions to this were *angiotensin-converting enzyme inhibitors* (OR per 10mmHg lower systolic blood pressure: 13.20; 95% CI: 2.14 to 81.24; p=0.005; rs4968783) and *potassium-sparing diuretics and aldosterone antagonists* (OR per 10mmHg lower systolic blood pressure: 0.17; 95% CI: 0.02 to 1.33; p=0.09; SNPs=3).

### Antihypertensive drug effect

We found little evidence for an overall effect of lowering systolic blood pressure on Alzheimer’s disease when combining all identified drug targets (OR per 10 mmHg lower systolic blood pressure: 1.14; 95% CI: 0.83 to 1.56; p=0.41; SNPs=59) (Supplementary Table 4).

### Systolic blood pressure effect

We also found little evidence for an overall effect of lowering systolic blood pressure on Alzheimer’s disease, without consideration of the associated drugs, as indicated by the OR of 1.04 (95% CI: 0.95 to 1.13; p=0.45; SNPs=135) per 10 mmHg lower systolic blood pressure (Supplementary Table 4).

### Sensitivity analyses

The Egger intercepts were close to zero for almost all analyses where they could be calculated (Supplementary Table 5). In addition, the estimates from the inverse variance weighted and MR-Egger methods were similar for all analyses with both the point estimate and confidence interval for the inverse variance weighted method almost contained within the confidence interval for the MR-Egger method (Supplementary Figure 3).

The analysis of individual targets identified some targets that were likely to be driving the drug class effects (Supplementary Figure 4). For example, the target *NR3C2* is estimated to be extremely protective (OR per 10 mmHg lower systolic blood pressure: 2.01e-3; 95% CI: 5.22e-6 to 0.78; p=0.04; rs71616586) and is likely to have contributed to the extremely protective effect observed for *potassium-sparing diuretics and aldosterone antagonists* (OR per 10 mmHg lower systolic blood pressure: 0.17; 95% CI: 0.02 to 1.33; p=0.09; SNPs=3).

## DISCUSSION

We found limited evidence to support an overall effect of lowering systolic blood pressure on Alzheimer’s disease risk (OR per 10 mmHg lower systolic blood pressure: 1.04; 95% CI: 0.95 to 1.13; p=0.45; SNPs=135). There was also limited evidence that lowering systolic blood pressure via antihypertensive drug classes affected Alzheimer’s disease. For example, calcium channel blockers had an OR of 1.53 (95% CI: 0.94 to 2.49; p=0.09; SNPs=17) and vasodilator antihypertensives had an OR of 0.98 (95% CI: 0.30 to 3.14; p=0.97; SNPs=11) per 10mmHg lower systolic blood pressure. This was reflected in the overall effect of lowering systolic blood pressure on Alzheimer’s disease when combining all identified drug targets, which had an OR of 1.14 (95% CI: 0.83 to 1.56; p=0.41; SNPs=59) per 10 mmHg lower systolic blood pressure. Despite this, we also report some extreme results, such as angiotensin-converting enzyme inhibitors, which were associated with an increased Alzheimer’s disease risk (OR per 10 mmHg lower systolic blood pressure: 13.29; 95% CI: 2.14 to 81.24; p=0.005; rs4968783).

The cause of these extreme results could be due to a competing mechanism, as illustrated in Figure 2. We estimated the effect of exposure to a given drug class on Alzheimer’s disease using the effect of the instrument for that drug class on both systolic blood pressure (instrument-exposure association) and Alzheimer’s disease (instrument-outcome association). Our analysis assumed that the effect we were estimating acted through systolic blood pressure, however there is potentially a competing mechanism by which the given drug class can affect Alzheimer’s disease. If a competing mechanism does exist and the instrument-exposure association (i.e. the effect of the drug class instrument on systolic blood pressure) is small, estimates from Mendelian randomization can become inflated as the competing mechanism means the instrument-outcome association (i.e. the effect of the drug class instrument on Alzheimer’s disease) remains large. This is more apparent if you consider the Wald ratio used to calculate the effect for single SNP instruments:

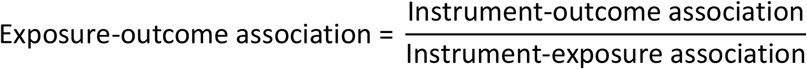

**Figure 2:**
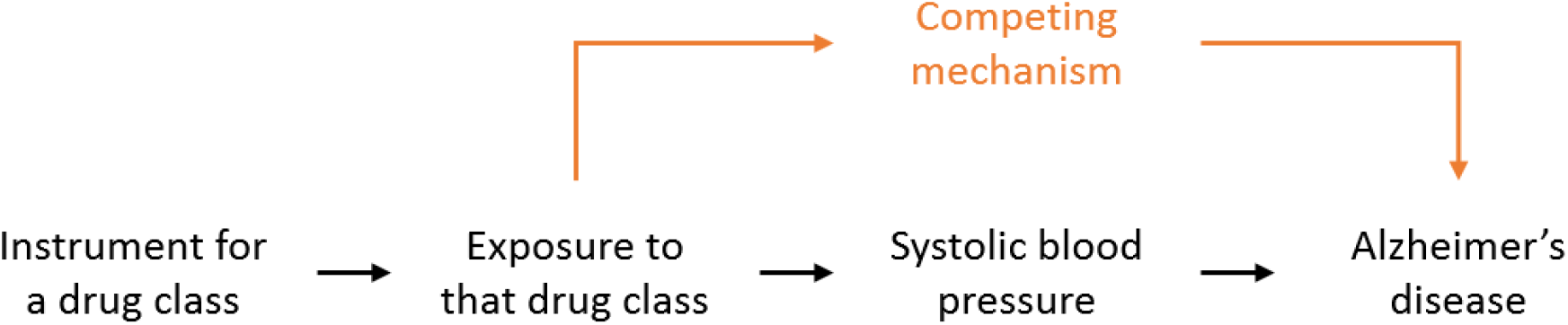
Mendelian randomization model in the presence of a competing mechanism

In our analysis, we found a small effect of systolic blood pressure on Alzheimer’s disease and our extreme results were for drug classes that may well act through competing mechanisms. For instance, returning to the example of *angiotensin-converting enzyme inhibitors*, angiotensin-converting enzyme is proposed to affect both vascular pathways (such as blood pressure) and have independent effects on amyloid beta. (15) In addition, *potassium-sparing diuretics and aldosterone antagonists*, which were also estimated to have an extreme effect (OR per 10 mmHg lower systolic blood pressure: 0.17; 95% CI: 0.02 to 1.33; p=0.09; SNPs=3), have previously been suggested to have a role, independent of blood pressure, in preventing cognitive decline. (33) This explanation for the extreme results observed for certain drug classes, along with the limited evidence for an effect among the remaining drug classes, indicates that antihypertensive drug classes are unlikely to have an effect on Alzheimer’s disease via systolic blood pressure.

### Comparison with existing literature

Two previous Mendelian randomization studies have studied the overall effect of systolic blood pressure on Alzheimer’s disease to date. These studies used different instruments and different systolic blood pressure GWAS, both to us and each other. (10,11) Østergaard et al found higher systolic blood pressure to be associated with a reduced risk of Alzheimer’s disease, while Larsson et al found little evidence of an effect of systolic or diastolic blood pressure with Alzheimer’s disease. Our results agree with Larsson et al in that there is unlikely to be an overall effect of systolic blood pressure on risk of Alzheimer’s disease. Gill et al recently conducted a study that combined MR using genetic variants related to antihypertensive targets with a PheWAS conducted in UK Biobank, however their analysis was restricted to beta-adrenoceptor blockers and calcium channel blockers. (12) Our results broadly agree with those reported by Gill et al for Alzheimer’s disease. There was a small overlap in the choice of SNPs used to instrument systolic blood pressure between our study and those previously reported however, there was very little overlap when considering our drug specific instruments (Supplementary Table 6). Using the previously reported instruments with our data, we were able to reproduce the previously reported results (Supplementary Figures 5 and 6).

Larsson et al recently conducted a systematic review and meta-analysis, which identified five randomized controlled trials that have investigated whether antihypertensives prevent dementia (not Alzheimer’s disease specifically). (4) Four of the five trials had point estimates that suggested a protective effect of antihypertensives compared to non-use, however three of these trials failed to exclude the null. This resulted in the meta-analysis finding an overall relative risk of 0.84 (95% CI: 0.69 to 1.02; p=0.10). It is worth highlighting that most studies described in the meta-analysis were from populations with high cardiovascular morbidity and were designed around cardiovascular related primary outcomes. In these trials, the proportion of dementia cases that derived from vascular mechanisms might be disproportionately high compared with other study populations. (34,35) This difference might explain the more favourable point estimate obtained in the meta-analysis. Since the publication of the meta-analysis, the first trial to consider an antihypertensive drug (*calcium channel blocker* Nilvadipine) as a direct intervention in Alzheimer’s disease has been published - it found no benefit of the treatment. (36)

### Strengths and limitations

A strength of our study was the use of two-sample Mendelian randomization that meant we were able to utilize the IGAP GWAS for our outcome data, which contains information on 17,008 Alzheimer’s disease cases and 37,154 controls. (9) The use of Mendelian randomization, over more conventional pharmacoepidemiological approaches, will have also addressed certain forms of confounding. This includes confounding by indication and confounding by the environmental and lifestyle factors of patients, which cannot be fully adjusted for using observational data. This is because measurement error and incomplete capture of all these potential confounding factors inevitably leads to residual confounding.

The limitations of this study included the risk of horizontal pleiotropy. We addressed this by conducting sensitivity analyses using MR-Egger when possible. Sensitivity analyses that considered the individual drug target effects also identified some heterogeneity that may have affected our drug class estimates – for example, the estimate for *potassium-sparing diuretics and aldosterone antagonist* may have seemed more protective due to the particularly large protective effect observed for one of the three targets under consideration: *NR3C2*. We were also limited by the fact that Mendelian randomization estimates the effect of lifelong exposure, while drugs typically have much shorter periods of exposure. This means that the effect sizes that we have estimated will not directly reflect what is observed in trials or clinical practice and may not distinguish critical periods of exposure. (37)

## CONCLUSION

This study helps to inform the growing knowledge around repurposing antihypertensive drugs for Alzheimer’s disease prevention by using a different method, subject to different biases, to assess this research question. We found little evidence to suggest that lowering systolic blood pressure itself will affect risk of developing Alzheimer’s disease. This was accompanied by limited evidence for many of the antihypertensive drug classes that we tested. This suggests that if specific antihypertensive drug classes do affect risk of Alzheimer’s disease, they are unlikely to do so via systolic blood pressure. Future research should consider this study, with other sources of evidence, in a triangulation framework to obtain a reliable answer concerning the potential repurposing of antihypertensives for Alzheimer’s disease prevention.

## Funding statement

This work was supported by the Perros Trust and the Integrative Epidemiology Unit. The Integrative Epidemiology Unit is supported by the Medical Research Council and the University of Bristol [grant number MC_UU_00011/1, MC_UU_00011/3].

## Competing interests declaration

Walker is currently working on a manuscript in collaboration with GlaxoSmithKline plc that explores whether Mendelian randomization can predict drug success but does not receive financial support from the company. Davies has worked on unrelated projects funded as part of the Global Research Awards For Nicotine Dependence, which is an independent grant giving body funded by Pfizer.

## Notes

#### Summary of Updates

Missing supplementary files.

