## Supplementary Figure for "Repurposing antihypertensive drugs for the prevention of Alzheimer’s disease: a Mendelian Randomization study"

### Supplementary Figures


1 Flow chart illustrating the instrument selection process. . . . . . . . . . . . . . . . . . . . . 2
2 Heat map of estimates for the effect of gene expression on systolic blood pressure. . . . 3
3 Estimates using MR-Egger. . . . . . . . . . . . . . . . . . . . . . . . . . . . . . . . . . 4
4 Target level estimates. . . . . . . . . . . . . . . . . . . . . . . . . . . . . . . . . . . . 5
5 Estimates using previously reported systolic blood pressure instruments. . . . . . . . . 6
6 Estimates using previously reported drug class instruments. . . . . . . . . . . . . . . . . 6

Supplementary Figure 1: Flow chart illustrating the instrument selection process.

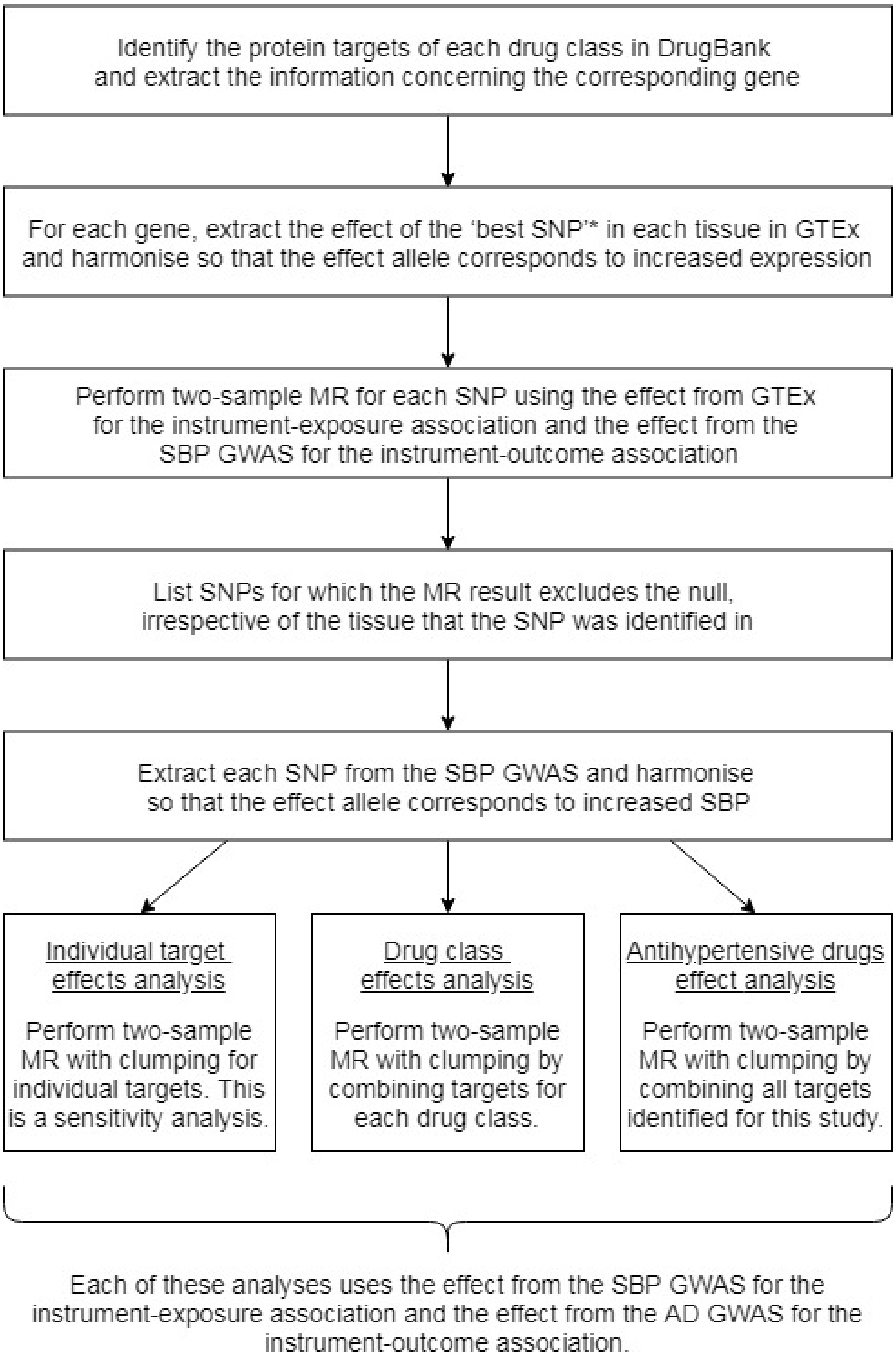


* The 'best SNP' is indciated by GTEx in the eGenes association file. It represents the variant with the smallest nominal p-value for a variant-gene pair.

Supplementary Figure 2: Heat map of estimates for the effect of gene expression on systolic blood pressure.

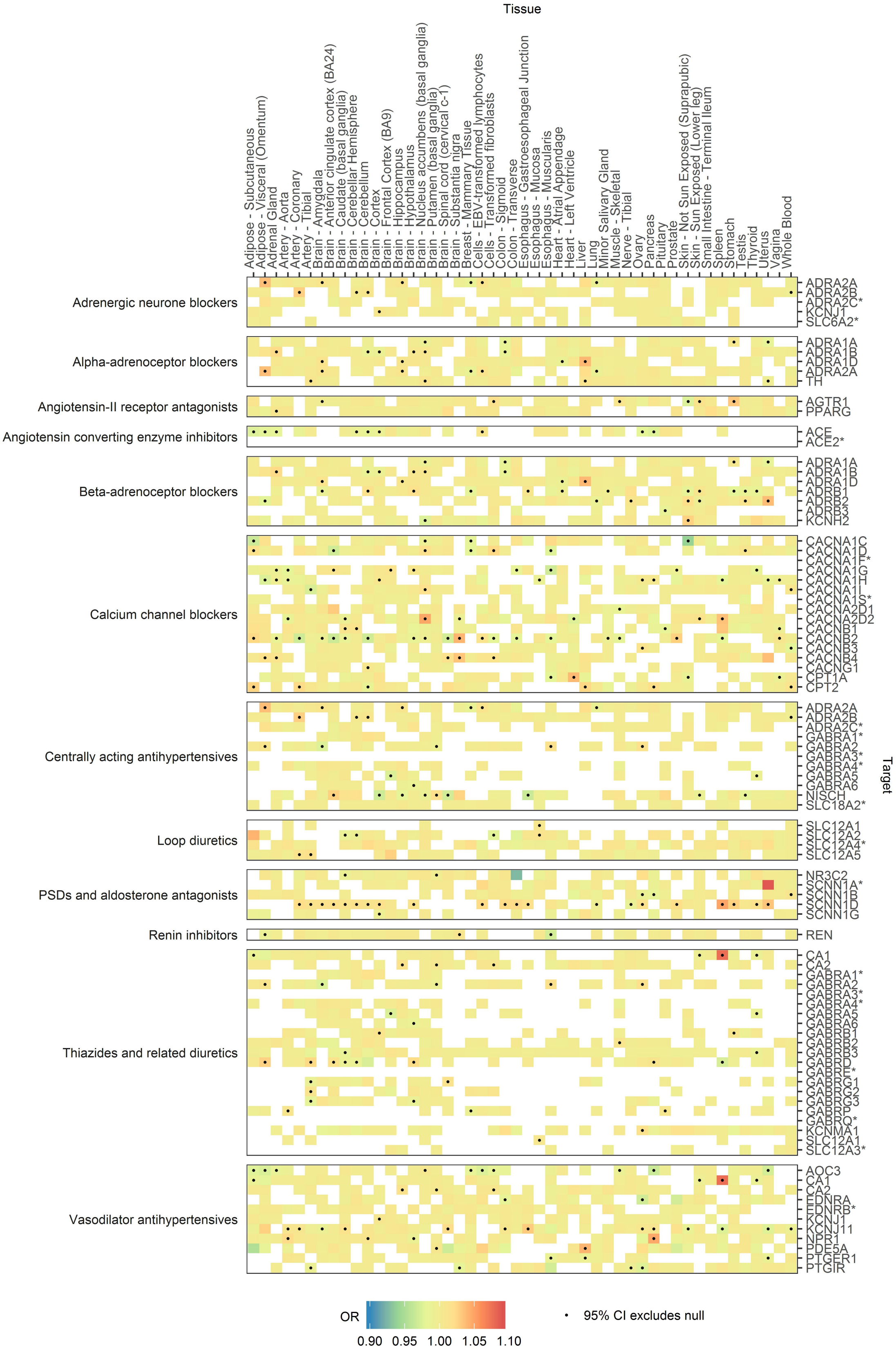


* No evidence for an effect on systolic blood pressure.

PSD: potassium sparing diuretic.

Supplementary Figure 3: Estimates using MR-Egger.

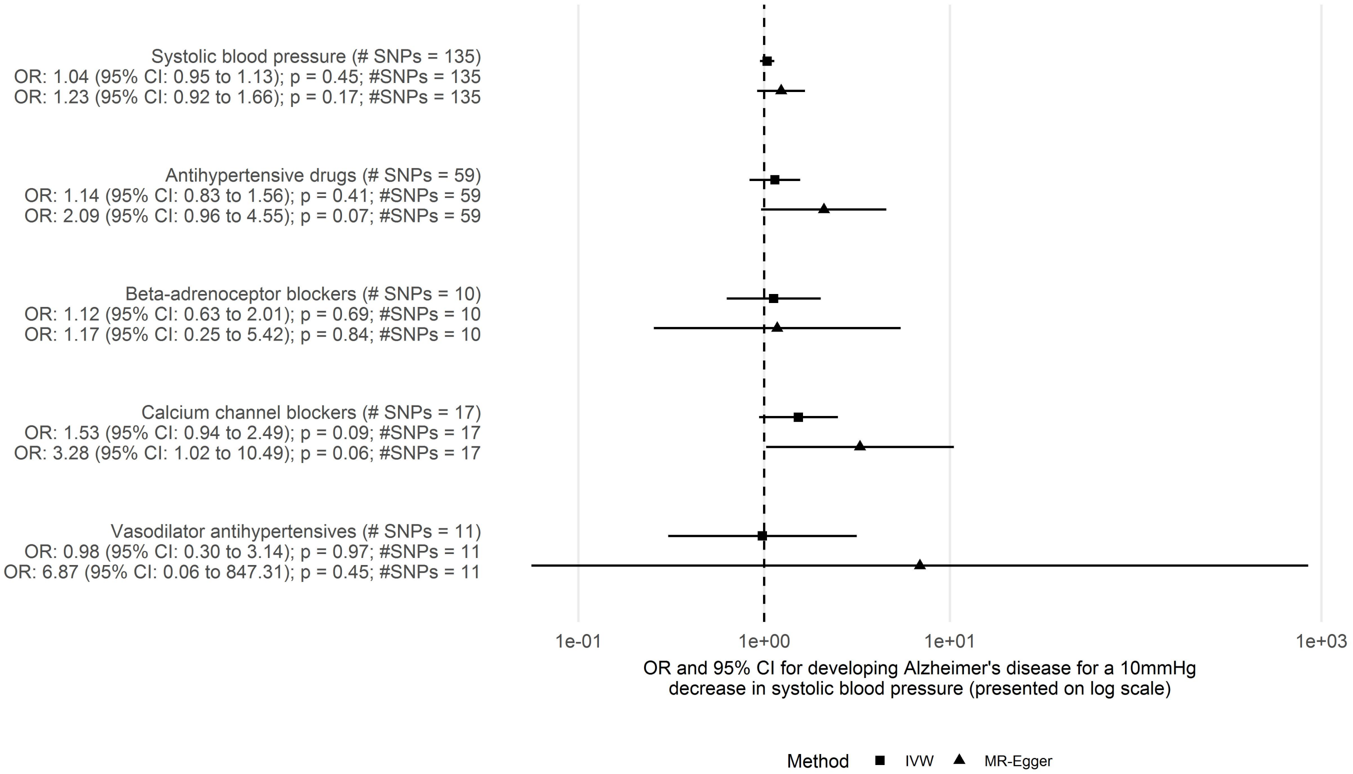


PSD: potassium sparing diuretic.

Supplementary Figure 4: Target level estimates.

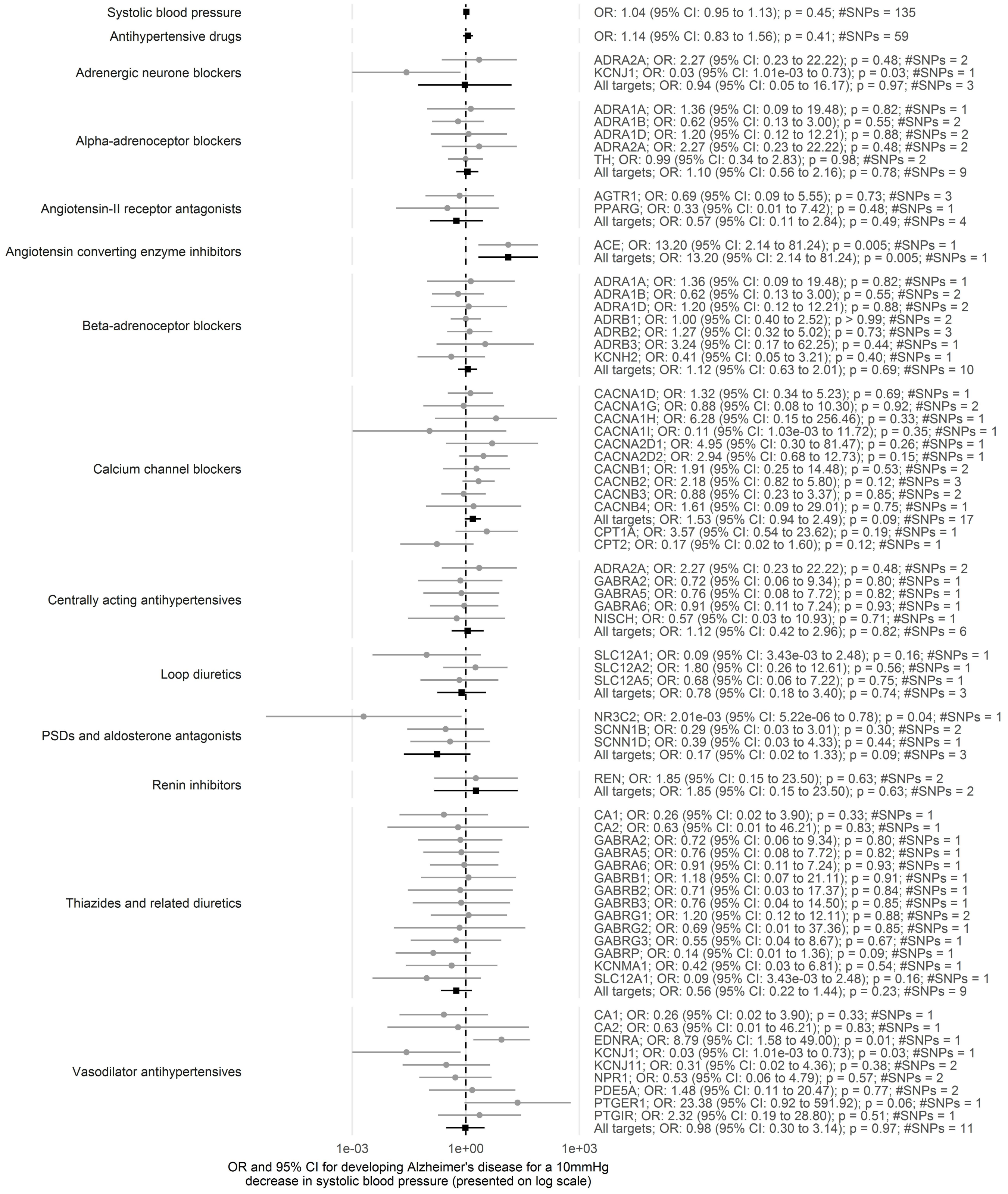


PSD: potassium sparing diuretic.

Supplementary Figure 5: Estimates using previously reported systolic blood pressure instruments.

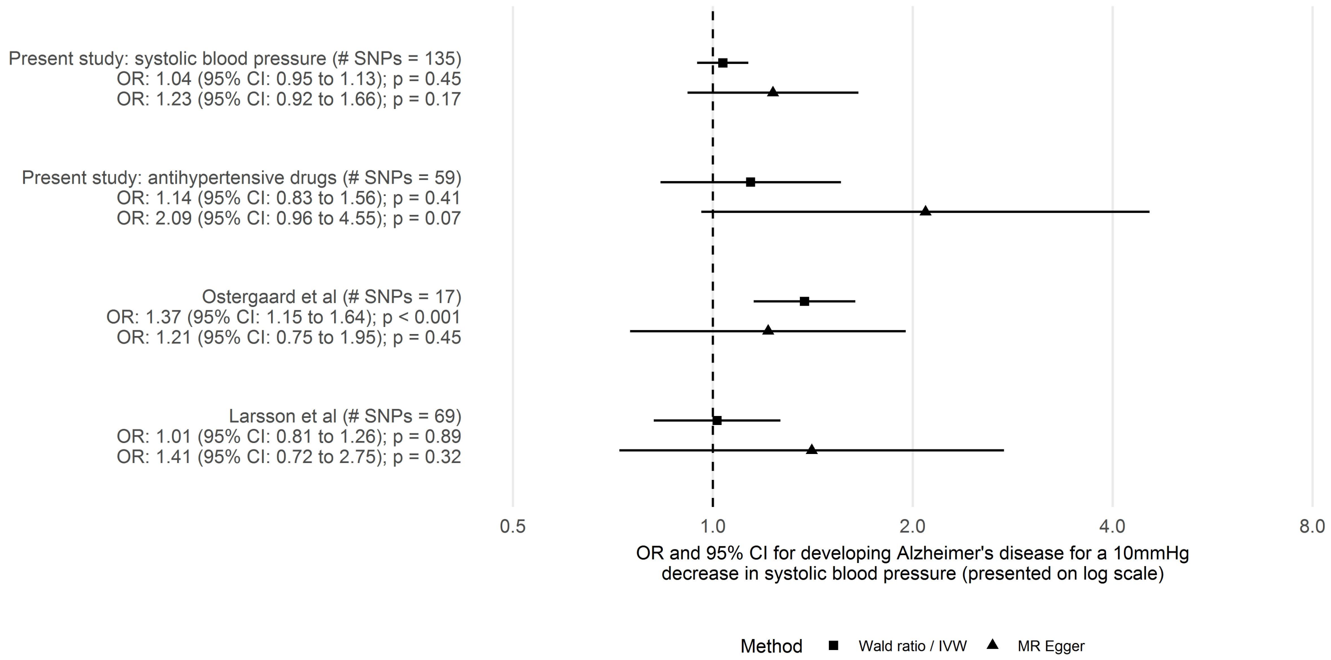


Note that we implemented our pipeline, including clumping, so there are a reduced number of SNPs in the instruments for Ostergaard et al (originally 25) and Larsson et al (originally 93) as our clumping criteria are likely to differ to those previously used.

Supplementary Figure 6: Estimates using previously reported drug class instruments.

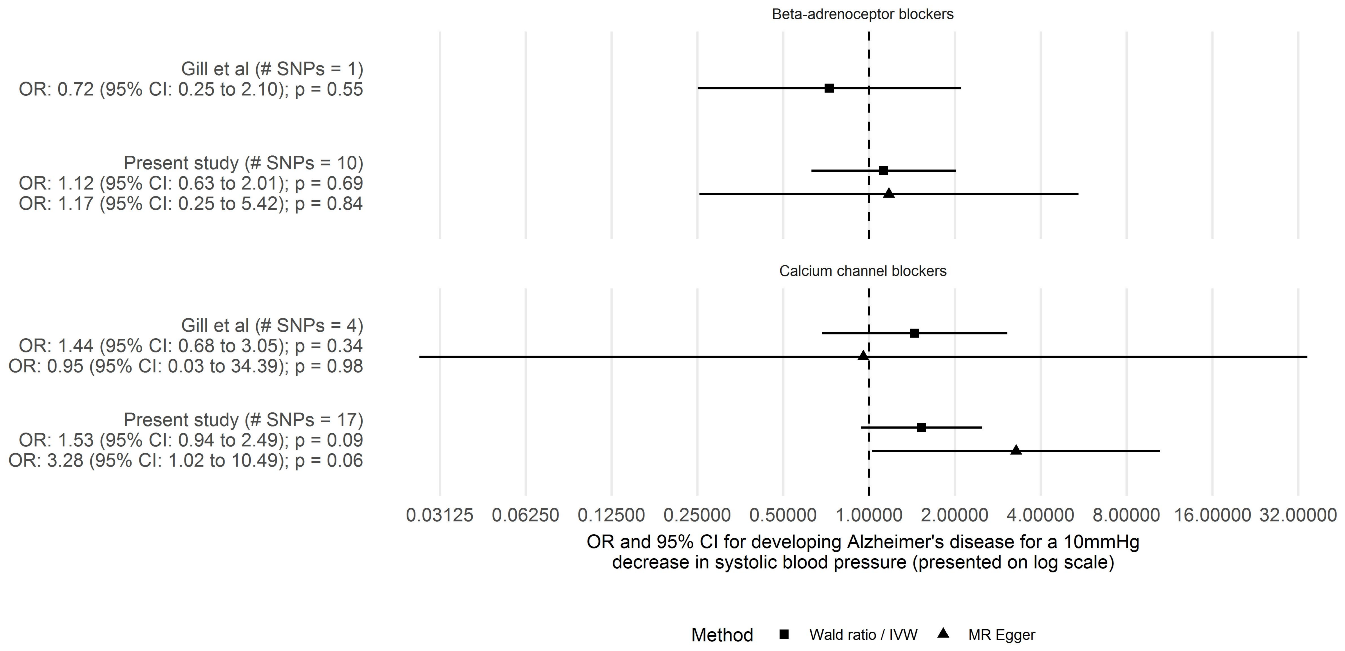


Note that we implemented our pipeline, including clumping, so there are a reduced number of SNPs in the instruments for beta-adrenoceptor blockers (originally 6) and calcium channel blockers (originally 24) obtained from Gill et al as our clumping criteria are likely to differ to those previously used.
